# Robust Bayesian Integrative Modeling of Single Cell and Spatially Resolved Transcriptomics Data

**DOI:** 10.1101/2025.04.22.650087

**Authors:** Huimin Li, Bencong Zhu, Xi Jiang, Ying Ma, Lin Xu, Qiwei Li

## Abstract

A recent technology breakthrough in spatially resolved transcriptomics (SRT) has enabled comprehensive molecular characterization at the cellular level while preserving spatial information. However, many SRT technologies (e.g., spatial transcriptomics) cannot achieve single-cell resolution. Instead, they measure the average gene expression of a mixture of cells within a spot arranged on a lattice. Here, we introduce iMOSCATO, a fully Bayesian model that integrates single-cell RNA sequencing (scRNA-seq) data and SRT data to simultaneously decompose cell-type compositions of spots and identify the underlying spatial domains. We incorporate the lattice structure by employing a Markov random field prior, improving the accuracy of both cell-type deconvolution and spatial domain detection. Moreover, we use a zero-inflated Dirichlet distribution to capture cell-type sparsity. Finally, iMOSCATO shows competitive performance in accuracy compared to existing methods in both simulation studies and two real data applications.

## 1. Introduction

Spatially resolved transcriptomics (SRT) is a collection of recently developed gene expression profiling technologies that provide biological information at the spot or cellular level while preserving the spatial organization of the tissue and its cellular microenvironment (Zhang et al., 2021; Marx, 2021; Li et al., 2023). Because of their enormous potential to reveal biological mechanisms through spatial analysis (De Bruin et al., 2014; Satija et al., 2015), SRT technologies are being increasingly applied across diverse fields, such as neuroscience, cancer research, and developmental biology, leading to an explosive growth in the generation of SRT data (Marx, 2021; Jung and Kim, 2023). Current SRT technologies are either imaging-based through single-molecule fluorescence *in situ* hybridization (FISH), such as seqFISH (Shah et al., 2018), seqFISH+(Eng et al., 2019), and STARmap (Wang et al., 2018), or next-generation sequencing-based (NGS-based) through spatial barcoding, such as spatial transcriptomics (ST) (Ståhl et al., 2016), 10x Visium (an improved ST platform), Slide-seq (Rodriques et al., 2019), and high-definition spatial transcriptomics (HDST) (Vickovic et al., 2019). Despite rapid technological advancements, most NGS-based SRT technologies still suffer from limited spatial resolution. Each spatial location may contain a mixture of cells belonging to potentially distinct cell types (Asp et al., 2020; Ståhl et al., 2016). As a result, NGS-based technologies effectively quantify the average gene expression level across all cells within that location. This resolution limitation constrains their ability to capture the gene expression data at single-cell resolution.

A central challenge in analyzing SRT data, therefore, is to quantify the proportion of different cell types within the regularly distributed spots, a process known as cell-type deconvolution. Cell-type deconvolution is an effective approach to improve the resolution of SRT data, enabling the discovery of spatial cell-type distributions and the characterization of complex tissue architecture (Liao et al., 2021; Ma and Zhou, 2022). Deconvolution of SRT data requires cell-type-specific gene expression information and tailored spatial methods. Consequently, current deconvolution methods integrate single-cell RNA-seq (scRNA-seq) data, which provides the necessary cell-type-specific gene expression profiles, with SRT data. However, most methods do not fully incorporate the rich spatial localization information in SRT data. Examples of such methods include the probabilistic-based methods SpatialDWLS (Dong and Yuan, 2021), RCTD (Cable et al., 2022), the non-negative matrix factorization-based (NMF-based) method SPOTlight (Elosua-Bayes et al., 2021), and the Bayesian-based method Cell2location (Kleshchevnikov et al., 2022) SpatialDWLS employs the weighted-least-squares approach to predict cell-type mixtures. RCTD models raw counts with Poisson distribution and applies supervised learning to decompose cell-type compositions. SPOTlight uses seeded NMF for cell-type deconvolution. Cell2location models raw counts with negative binomial distributions and uses the gene expression signature of the cell subpopulations in scRNA-seq data to estimate cell-type compositions.

Spatial localization information in SRT captures the relative distances between tissue locations and provides potentially valuable insights for deconvolution. Hematoxylin and eosin (H&E) staining images accompanying SRT datasets (Ståhl et al., 2016) highlight the spatial segregation of cell types and the similarity in cell-type composition between neighboring regions. Neighboring tissue locations tend to have more similar cell-type compositions than distant ones, so modeling the neighborhood similarity and spatial correlations can help borrow composition information across the tissue, improving deconvolution accuracy. Some state-of-the-art methods incorporate the spatial localization information. For example, CARD (Ma and Zhou, 2022) is an NMF-based model that captures the spatial correlation structure using a conditional autoregressive (CAR) modeling assumption. Despite the advantages of these deconvolution methods, they have several limitations. First, SRT data often contain a substantial proportion of zeros (e.g., 60% to 99%), which can significantly reduce statistical power; this issue is not addressed by any of the aforementioned methods. Second, each spot in NGS-based SRT data may contain a varying number of cells, ranging from just a few to hundreds (Saiselet et al., 2020; Moncada et al., 2020), leading to a cell-type sparsity issue. However, none of the methods mentioned above address this challenge. Lastly, most methods overlook the rich spatial localization information available in SRT data, which can reduce deconvolution accuracy, as discussed earlier.

Another challenge in analyzing SRT data is to detect clinically or biologically meaningful spatial domains by partitioning regions with similar molecular or morphological characteristics. Spatial domain detection is crucial for downstream analyses, such as domain-based differential expression analysis (Thrane et al., 2018), trajectory analysis (Shang and Zhou, 2022), and functional pathway analysis (Wang et al., 2024). Recent advancements in SRT clustering approaches can be classified into two categories based on the additional information that they use to improve spatial domain detection performance. The first group incorporates the geospatial profiles by employing Markov random field (MRF) model to account for spatial correlations, as seen in methods such as BayesSpace (Zhao et al., 2021), SC-MEB (Yang et al., 2022), DR-SC (Liu et al., 2022), BayesCafe (Li et al., 2024), and BNPSpace (Zhu et al., 2023). The second group leverages histology images available in SRT data. For example, stLearn (Pham et al., 2020) extracts tissue morphological features, SpaGCN (Hu et al., 2021) utilizes RGB color values from histology images, and iIMPACT (Jiang et al., 2023) incorporates detailed morphological information from AI-reconstructed histology images. Although the aforementioned methods have shown their strengths in detecting spatial domains, they also come with clear limitations. First, these methods often transform count data, which typically contains a large proportion of zeros, into continuous data for more convenient statistical modeling. However, this transformation may not accurately capture the underlying data generation mechanism and can cause information loss (Sun et al., 2020). Second, the high sparsity existing in SRT data may significantly reduce statistical power and is not handled by most of the aforementioned methods. Table S2 provides a brief summary of the two real datasets analyzed in this paper, both of which exhibit high sparsity. Lastly, none of the existing methods can tackle these two important challenges simultaneously, even though they are closely intertwined in SRT data. Effectively addressing both is essential for accurately capturing the complex spatial organization and underlying cell-type heterogeneity, both of which drive meaningful biological insights. A joint solution can better preserve spatial context while enhancing the resolution and interpretability of the transcriptomic signals.

To overcome the aforementioned limitations, we developed iMOSCATO, a fully Bayesian model that integrates scRNA-seq and SRT data for deconvolving and clustering SRT data. iMOSCATO directly models the molecular profiles of SRT data using a zero-inflated negative binomial (ZINB) mixture model to account for the zero-inflation and over-dispersion observed in SRT data while avoiding the need for an *ad hoc* data normalization method. Additionally, iMOSCATO leverages a zero-inflated Dirichlet distribution (ZIDD) to capture cell-type sparsity in cell-type compositions, enabling simultaneous deconvolution and spatial domain detection. Furthermore, iMOSCATO employs an MRF prior to integrate the geospatial profiles of SRT data, improving clustering accuracy. We demonstrated the advantages of iMOSCATO through a comprehensive simulation study that explores various cell-type sparsity and zero-inflation scenarios. We also applied iMOSCATO to two NGS-based SRT datasets, showing improved deconvolution and clustering accuracy compared to existing methods.

## 2. Model

In this section, we introduce iMOSCATO for cell-type deconvolution and spatial domain detection. The overall workflow is depicted in Figure 1. Figure S1 and Table S1 summarize the graphical and hierarchical formulation of the proposed model in the supplementary materials.

**Figure 1:**
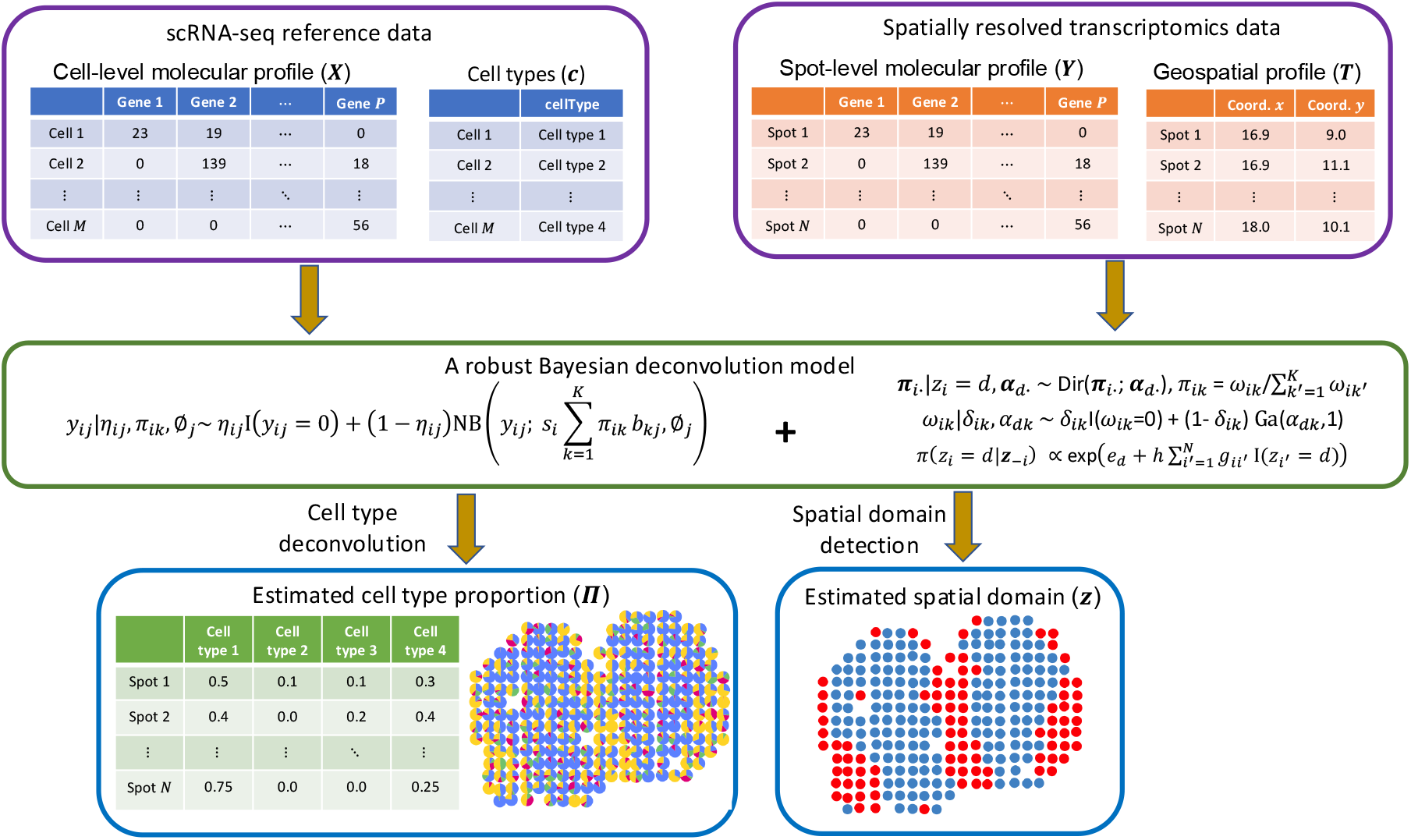
The schematic diagram of the proposed iMOSCATO model.

### 2.1 Data Preparation

Before introducing the main components, we summarize the observed data as follows. Let an *M* -by-*P*^*x*^ count matrix ***X***^Full^ denote the cell-level molecular profiles generated by a single-cell RNA sequencing (scRNA-seq) technique, where each entry *x*_*mj*_ ∈ ℕ, *m* = 1, …, *M, j* = 1, …, *P, P* + 1, · · ·, *P*^*x*^ is the read count observed in cell *m* (a cell with known cell type) for gene *j*. Let an *M* -dimensional vector ***c*** = (*c*_1_, …, *c*_*M*_)^⊤^ to allocate the *M* cells into *K* mutually exclusive groups (i.e. *K* cell types), with *c*_*m*_ = *k, k* = 1, …, *K* indicating that cell *m* belongs to cell type *k*.

Let a *N* -by-*P*^*y*^ count matrix ***Y*** ^Full^ denote the spot-level molecular profiles generated by an ST technique, where each entry *y*_*ij*_ ∈ ℕ, *i* = 1, · · ·, *N, j* = 1, · · ·, *P, P* + 1,, *P*^*y*^, is the read count observed in spot *i* for gene *j*. Let a *N* -by-2 matrix ***T*** represent the geospatial profiles, where each row ***t***_*i*·_ = (*t*_*i*1_, *t*_*i*2_) ∈ ℝ^2^ gives the coordinates of spot *i* in a compact subset of the two-dimensional Cartesian plane (see examples in Figure 1).

We assume the high-dimensional spaces are identical for the scRNA-seq and SRT data. This could be done by fully intersecting the sets of scRNA-seq and ST gene, then results in a *M* -by-*P* scRNA-seq count matrix ***X*** and a *N* -by-*P* ST count matrix ***Y*** analyzed in the model. We use ***y***_*i*·_ = (*y*_*i*1_, …, *y*_*iP*_) and ***y***_·*j*_ = (*y*_1*j*_, …, *y*_*Nj*_)^⊤^ to denote the *i*th row and *j*th column of the count matrix ***Y***.

Let ***v*** = (*v*_1_, · · ·, *v*_*M*_)^⊤^ and ***s*** = (*s*_1_, …, *s*_*N*_)^⊤^ be the collection of the sample-specific size factor of scRNA-seq data and SRT data, respectively, which reflects many nuisance effects across sample points, including, but not limited to: 1) reverse transcription efficiency; 2) amplification and dilution efficiency; 3) sequencing depth. To ensure identifiability between *v*_*m*_, *s*_*i*_ and normalized expression levels, we follow (Sun et al., 2020) to set *v*_*m*_ and *s*_*i*_ proportional to the summation of the total number of read counts across all genes observed at cell *m* and spot *i*, respectively, i.e., 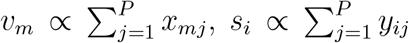 combined with a constraint of 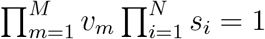. It results in 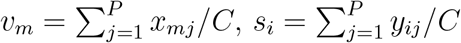, where 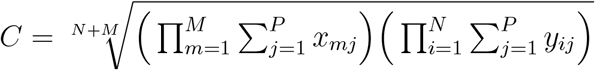.

### 2.2 Modeling the SRT Molecular Profiles via a ZINB Model

Previous studies on scRNA-seq data analysis have suggested that accounting for the large proportion of zeros in the model can lead to a substantial improvement in model fitting and accuracy of identifying differentially-expressed genes (Finak et al., 2015; Lun et al., 2016). Therefore, we start by considering a ZINB model to model the read counts of SRT data:

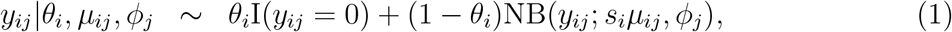

where we use I(·) to denote the indicator function and NB(*µ, ϕ*), *µ, ϕ >* 0 to denote a negative binomial (NB) distribution with expectation *µ* and dispersion 1*/ϕ*. With this NB parameterization, the probability mass function (p.m.f) is written as 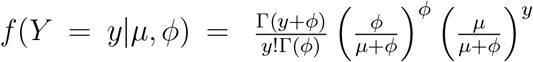, with variance Var(*Y*) = *µ* + *µ*^2^*/ϕ*, thus allowing for over-dispersion. A small value of *ϕ* indicates a large variance-to-mean ratio, while a large value approaching infinity simplifies the NB model to a Poisson model with the same mean and variance. In this model, we constrain one component of the mixture model to degenerate at zero, thereby allowing for zero-inflation. The sample-specific parameter *θ*_*i*_ ∈ (0, 1) can be treated as the proportion of extra zero (i.e., false zero or structural zero) counts at spot *i*.

The mean in the NB distribution is further decomposed into two multiplicative effects: the size factor *s*_*i*_ ∈ ℝ^+^ and the normalized expression level *µ*_*ij*_ ∈ ℝ ^+^. After adjusting for the global sample-specific effect, *µ*_*ij*_ can be regarded as the normalized expression level of gene *j* observed at spot *i*. After accounting for zero-inflation (*via θ*_*i*_), over-dispersion (*via ϕ*_*j*_), and sample point heterogeneity (*via s*_*i*_), our modeling approach produces a denoised version of gene expression levels, represented by *µ*_*ij*_.

To facilitate model fitting, we rewrite Equation (1) by introducing a latent indicator variable

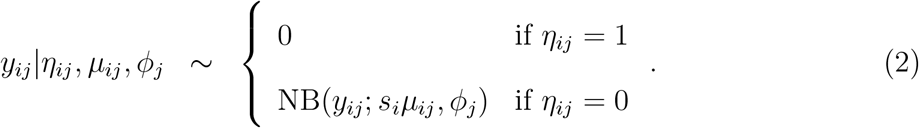

We impose an independent Bernoulli prior on *η*_*ij*_, i.e. *η*_*ij*_ ∼ Bern(*θ*_*i*_). Setting *θ*_*i*_ ≡ *θ* = 0.5 results in a weakly informative setting. We assume a gamma prior for all dispersion parameters *ϕ*_*j*_, i.e. *ϕ*_*j*_ ∼ Ga(*a*_*ϕ*_, *b*_*ϕ*_) and suggest small values such as *a*_*ϕ*_ = *b*_*ϕ*_ = 0.001 for a weakly informative setting.

### 2.3 Deconvolving Cell Types and Detecting Spatial Domains via a ZIDD Model

Cell-type deconvolution aims to identify cell types and their relative proportions contributing to a spot. Without loss of generality, we assume that the sample-specific normalized expression level equals to the weighted average of normalized gene expressions from each cell type within a spot, given by 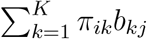, where *π*_*ik*_ ∈ [0, 1] is the underlying proportion of cells with cell type *k* within spot *i*. ***π***_*i*·_ = (*π*_*i*1_, …, *π*_*iK*_)^⊤^ with the constraint 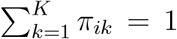 for all *i* = 1, · · ·, *N*. *b*_*kj*_ denotes the normalized expression level of gene *j* for cell type *k*. Following CARD (Ma and Zhou, 2022), we derive *b*_*kj*_ from the scRNA-seq reference data. Conditioning on *η*_*ij*_ = 0, we can rewrite our model as follows:

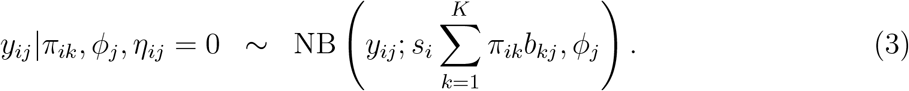

If *η*_*ij*_ = 1, we assume *y*_*ij*_ = 0 irrespective of *π*_*ik*_ and *ϕ*_*j*_.

To detect spatial domains, we assume that the *N* spots belong to different spatial domains *D*. To model cell type sparsity, we introduce a zero-inflated Dirichlet distribution (ZIDD). We place a Dirichlet prior on ***π***_*i*·_, assuming each spatial domain is characterized by a unique concentration parameter vector ***α***_*d*_·. Thus,

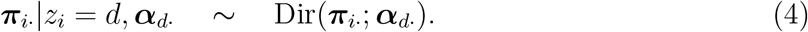

where ***z*** = (*z*_1_, · · ·, *z*_*N*_)^⊤^ is the spatial domain allocation vector, *z*_*i*_ = *d* if the *i*th spot belongs to the spatial domain *d* for *d* = 1, 2 · · ·, *D*, where *D* is the number of spatial domains. ***α***_*d*·_ = (*α*_*d*1_, · · ·, *α*_*dK*_)^⊤^ is the collection of concentration parameters in spatial domain *d*. We assume a gamma prior for *α*_*dk*_, that is, *α*_*dk*_ ∼ Ga(*a*_*α*_, *b*_*α*_) and recommend small values such as *a*_*α*_ = *b*_*α*_ = 0.001 for a weakly informative setting.

The number of spatial domains, *D*, can be determined based on prior biological knowledge when available. In the absence of such information, the modified Bayesian Information Criterion (mBIC) (Wang et al., 2007) can be used as a criterion to select K.

The Dirichlet distributed underlying proportion ***π***_*i*·_ can be equivalently represented as normalized gamma random variables 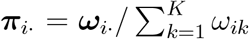, where ***ω***_*i*·_ = (*ω*_*i*1_, · · ·, *ω*_*iK*_)^⊤^ are unnormalized random variables and *ω*_*ik*_|*α*_*ik*_ ∼ Ga(*α*_*ik*_, 1). Since spots may not contain all cell types (Koslovsky, 2023), we adopt a zero-inflated Gamma distribution for *ω*_*ik*_ to model the potential zero-inflation, leveraging the data augmentation technique mentioned above,

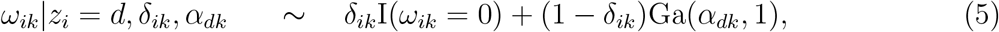

where the indicator variable *δ*_*ik*_ ∈ {0, 1} indicates the presence of cell type *k* in spot *i*. if *δ*_*ik*_ = 1, *ω*_*ik*_ = *π*_*ik*_ = 0, indicating that spot *i* does not contain cell type *k*, if *δ*_*ik*_ = 0, then *π*_*ik*_ *>* 0. We impose an independent Bernoulli prior for *δ*_*ik*_, i.e., *δ*_*ik*_ ∼ Bern(*q*_*i*_). Setting *q*_*i*_ ≡ *q* = 0.5 results in a weakly informative setting.

### 2.4 Integrating the ST Geospatial Profiles via an MRF Prior Model

To efficiently incorporate available spatial information, we impose a Markov random field (MRF) prior on ***z*** to encourage neighboring spots to cluster into the same group:

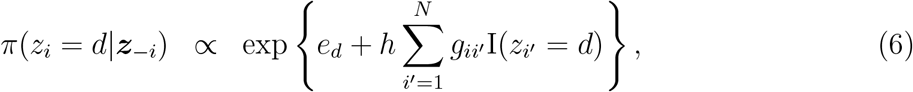

where *e*_*d*_ and *h* are hyperparameters to be chosen and ***z***_−*i*_ denotes the vector of ***z*** excluding the *i*th element. Here *e*_*d*_ controls the abundance of each domain, and *h* controls the strength of spatial dependency. We can also write the joint MRF prior on ***z*** by

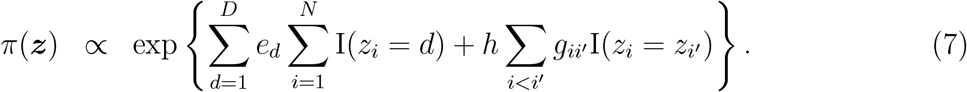

As a result, neighboring locations are more likely to be assigned to the same domain. The adjacency matrix ***G***, an *N* × *N* symmetric matrix, can be constructed based on the geospatial profiles ***T*** to define the neighborhood structure, with *g*_*ii*_*′* = 1 if spots *i* and *i*^′^ are neighbors, and *g*_*ii*_*′* = 0 otherwise. For ST and 10x Visium platforms, ***G*** is created based on their square and triangular lattices, respectively. For other SRT technologies, ***G*** is constructed using a Voronoi diagram (Okabe et al., 2009). Note that if a sample point does not have any neighbors, its prior distribution reduces to a multinomial prior with parameter ***q*** = (*q*_1_, …, *q*_*D*_)^⊤^ where 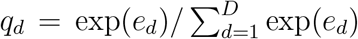 is a multinomial logistic transformation of *e*_*d*_. Although the parameterization is somewhat arbitrary, a careful selection of *h* is crucial. In particular, a large value of *h* may lead to a phase transition problem (i.e., all sample points are assigned into the same cluster). We discuss this issue in our paper (Li et al., 2024) and follow it by setting *e*_1_ = · · · = *e*_*D*_ = 1 and *h* = 1.

## 3. Model Fitting

### 3.1 MCMC Algorithm

We employ a Markov chain Monte Carlo (MCMC) algorithm to update all parameters in our model. Our model simultaneously infers cell-type mixtures across spots *via* **Π** and partitions spots into distinct spatial domains *via* ***z***. We jointly update the cell type proportions *π*_*ik*_ and their extra zero indicators *δ*_*ik*_ *via* an *add-delete* algorithm. The domain labels *z*_*i*_ and false zero indicators *η*_*ij*_ are updated using a Gibbs sampler, while the remaining parameters are updated using the random walk Metropolis-Hastings (RWMH) algorithm. We note that this algorithm is sufficient to guarantee ergodicity for our model. Full details can be found in the supplementary materials.

### 3.2 Posterior Inference

Our primary goal is to predict the cell-type compositions of the spots through the matrix **Π** and to identify spatial domains through the vector ***z***. We obtain posterior inference on these parameters by post-processing the MCMC samples after burn-in. We obtain posterior inference on these parameters by postprocessing the MCMC samples after burn-in. We summarized the posterior distributions of **Π** and ***z*** by using the *maximum a posteriori* (MAP) estimates, which corresponds to the configuration with highest conditional posterior probability among those drawn by the MCMC sampler. Specifically, we define 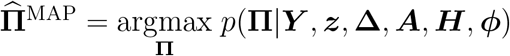 and 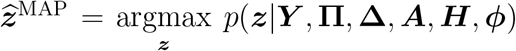. In addition, we can obtain a summary of ***z*** based on the pairwise probability matrix (PPM). The PPM is an *N* -by-*N* symmetric matrix whose elements are the posterior pairwise probabilities of co-clustering, that is, the probability that spot *i* and spot *i*^′^ belong to the same domain: 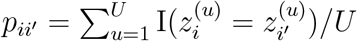 (Dahl, 2006), *where u* = 1, · · ·, *U* indicates the MCMC iterations after burn-in. Then, the point estimate of the clustering, 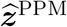, can be obtained by minimizing the sum of squared deviations of its association matrix from the PPM:

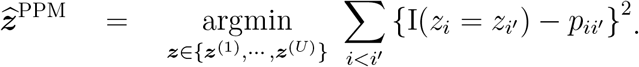

The PPM estimate has the advantage of utilizing information from all clusterings through the PPM. It is also intuitively appealing because it selects the “average” clustering rather than forming a clustering *via* an external, *ad hoc* clustering algorithm.

## 4. Simulation Study

In this section, we briefly summarize the simulation study. A detailed description is available in Web Appendix B. We followed the data generative schemes described in (Li et al., 2024), based on the spatial pattern constructed from a mouse olfactory bulb (MOB) study, as shown in Figure 2. The MOB pattern contains *N* = 260 spots and *D* = 2 spatial domains. We simulated *N* = 500 cells with *K* = 4 cell types and *p* = 1, 000 genes, including *p*_*γ*_ = 20 cell-type marker genes. We simulated cell-level count *x*_*mj*_ and spot-level count *y*_*ij*_ using a ZINB mixture model and generated cell type proportions *π*_*ik*_ from a zero-inflated Gamma distribution. The data generative model is detailed in Web Appendix B.1 in the supplementary materials, while the prior choice and MCMC algorithm settings are presented in Web Appendix B.2.

**Figure 2:**
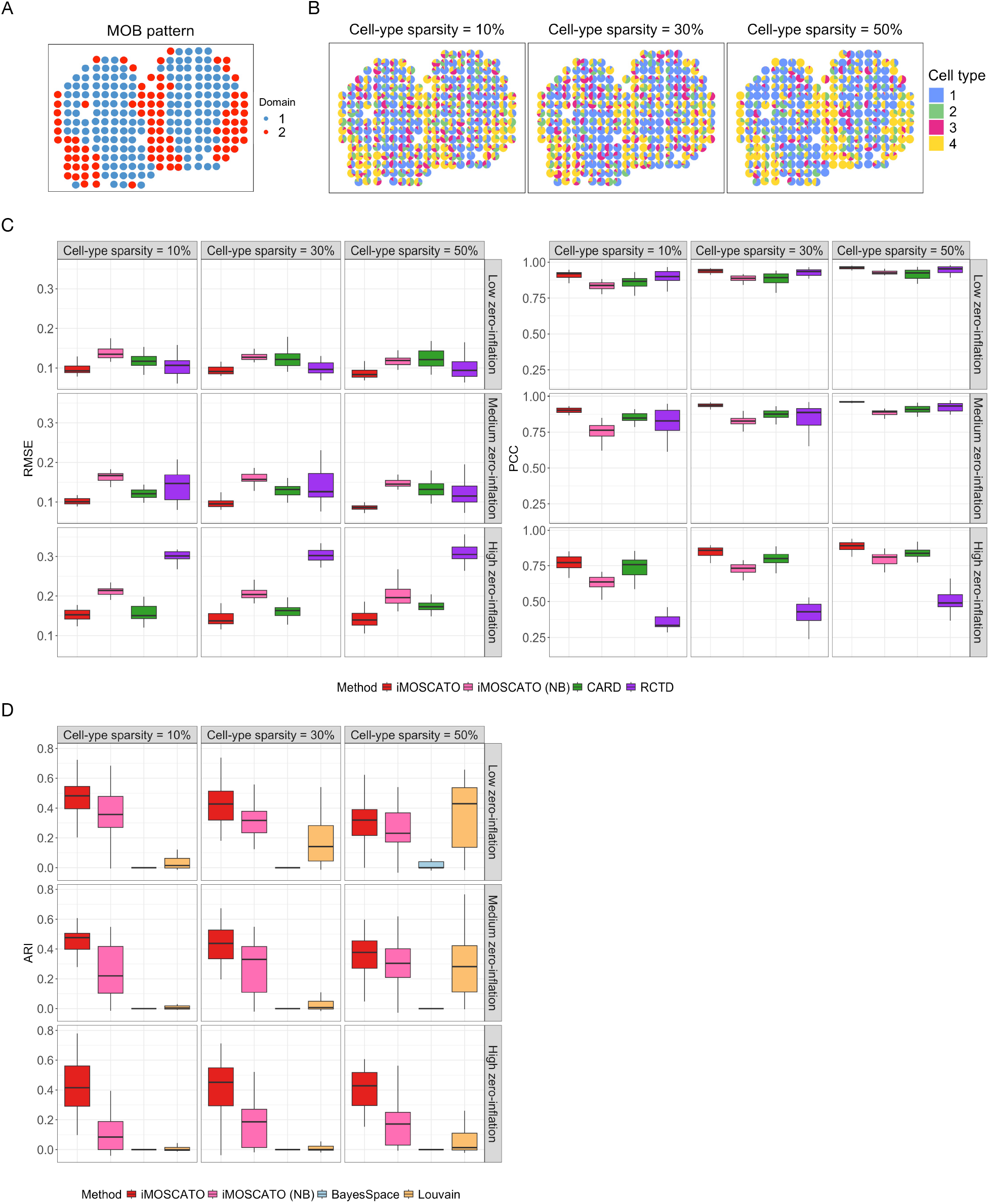
The simulation study. A. The spatial pattern used to generate the simulated data, which was constructed from the mouse olfactory bulb (MOB) study. B. Examples of simulated cell-type composition patterns across different cell-type sparsity settings. C. The boxplots of root mean square errors (MSEs) and Pearson correlation coefficients (PCCs) achieved by different cell-type deconvolution methods under different scenarios in terms of count zero-inflation and cell-type sparsity settings. D. The boxplots of adjusted Rand indices (ARIs) achieved by different spatial domain detection methods under different scenarios in terms of count zero-inflation and cell-type sparsity settings.

We followed Ma and Zhou (2022) to evaluate the deconvolution performance by calculating the root mean square error (RMSE), given by 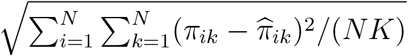, and Pearson correlation coefficient (PCC) between the estimated cell-type composition and the underlying truth at each spatial location. RMSE assesses the exact difference between the estimated cell-type composition and the ground truth, while the PCC is used to evaluate the similarity between these two components. To assess the performance of spatial domain detection, we used the adjusted Rand index (ARI) (Hubert and Arabie, 1985), which is a corrected-for-chance version of the Rand index (Rand, 1971). The Rand index measures the similarity between two partitions, with values ranging from 0 to 1. Larger ARI values indicate better performance, and a value of 1 indicates a perfect agreement between the two partitions.

For the deconvolution task, the competing methods we consider are CARD and RCTD. For the spot clustering task, the competing methods are BayesSpace and Louvain. To demonstrate the contribution of the zero-inflated part of iMOSCATO, we implemented a negative binomial version of iMOSCATO (i.e. fix *θ*_*i*_ = 0 ∀*i*), named iMOSCATO (NB).

Figure 2C displays the boxplots of RMSEs and PCCs achieved by different deconvolution methods over 30 replicates in the nine scenarios, where each subplot represents the result of the combination of cell-type sparsity levels and zero-inflation levels. In all 9 scenarios, iMOSCATO achieved the smallest RMSE and highest PCC. As the zero-inflation level increases, the performances of iMOSCATO, iMOSCATO (NB), and CARD are robust but RCTD decreases dramatically, indicating that RCTD is sensitive to the dropout events in the RNAseq data. In contrast, all other methods, including iMOSCATO (NB) suffered from excess zeros, suggesting that the variation caused by an excess of zero counts was not properly addressed. As a supplement, we also evaluated the agreement between the spatial domain ground truth and dominant cell type. iMOSCATO achieved highest value of ARIs, followed by iMOSCATO (NB) and CARD (see Figure S2A). We also observed that iMOSCATO demonstrated superior performance in the presence of high cell-type sparsity, accurately identifying the non-existing cell types with high area under the curve (AUC) values, as shown in Figure S2B. This result strongly supports the advantage of iMOSCATO in modeling cell-type sparsity through the ZIDD model.

Figure 2D displays the boxplots of ARIs achieved by different spatial domain detection methods over 30 replicates under the nine scenarios.

BayesCafe was designed to detect spatial domains and identify discriminating genes in SRT data. Since our simulation study generated cell-type marker genes for scRNA-seq data, we did not include BayesCafe in the performance comparison. For spatial domain identification, iMOSCATO achieved the best performance across all settings, followed by iMOSCATO (NB) and Louvain. In contrast, BayesSpace often failed to discriminate two domains, resulting in ARI values close to zero. iMOSCATO also maintained robust performance with respect to increasing zero inflation levels, while the performance of other methods declined significantly. This result highlights the contributions of the zero-inflated model for the count RNA sequencing data. Overall, the simulation study clearly demonstrated that the joint modeling of cell-type deconvolution and spatial domain detection through iMOSCATO enhances the performance of both tasks.

## 5. Real Data Analysis

iMOSCATO was applied to two real datasets, and the results are reported below. For an overview, Table S4 in the supplementary materials summarizes the deconvolution and clustering accuracy along with computational efficiency. Additionally, we employed the Normalized Mutual Information (NMI) (Strehl and Ghosh, 2002), a well-established metric in machine learning and data mining (Knops et al., 2006; Do et al., 2021; Molaei et al., 2021). This metric complements the ARI by addressing its potential sensitivities to factors such as cluster size and shape.

### 5.1 Application to the Mouse Olfactory Bulb ST Data

To further assess iMOSCATO’s performance, we first applied it to a publicly available SRT dataset from a mouse olfactory bulb (MOB) study (Ståhl et al., 2016). It is accessible on the website of the Spatial Research Lab at the KTH Royal Institute of Technology (https://www.spatialresearch.org/). We followed (Ma and Zhou, 2022) to focus on the MOB replicate 12, which contains 16,034 genes measured on 282 spots. The MOB data includes four main anatomic layers organized in an inside–out fashion, annotated by CARD (Ma and Zhou, 2022) based on histology: the granule cell layer (GCL), the mitral cell layer (MCL), the glomerular layer (GL) and the nerve layer (ONL). We used scRNA-seq data (Tepe et al., 2018) from Gene Expression Omnibus (GEO; accession number GSE121891) on the same tissue for deconvolution. scRNA-seq data contains 18,560 genes and 12,801 cells, corresponding to 5 cell types: external plexiform layer interneurons (EPL-IN), granule cells (GC), mitral and tufted cells (M-TC), olfactory sensory neurons (OSNs), and periglomerular cells (PGC). The cell type information in scRAN-seq data and spatial domain information in SRT data is presented in Table S3. We filtered out spots with fewer than 100 total counts across all genes (Ma and Zhou, 2022; Li et al., 2024) and genes with more than 90% zero read counts on all spots (Li et al., 2021, 2024). This quality control (QC) procedure led to a final set of *n* = 278 spots and 9, 904 genes in SRT data. In the scRNA-seq data, we filtered out genes that have zero counts on all cells and filtered out cells that have zero counts on all genes. These filtering criteria led to a final set of 17,812 genes and 12,801 cells for analysis. We evaluated the convergence of the MCMC algorithm based on estimated cell-type proportions for all four chains and found that their pairwise Pearson correlation coefficients ranged from 0.93 to 0.95 (see Figure S3A), along with trace plot of one estimated cell-type proportion in Figure S3B, indicating good MCMC convergence. We then aggregated the results of all four chains. We compared the deconvolution results of iMOSCATO with those of iMOSCATO (NB), CARD, and RCTD, using their default settings and set the number of domains to four (*D* = 4) for iMOSCATO and iMOSCATO (NB). As shown in Figure 3A, the cell-type compositions inferred by iMOSCATO accurately captured the expected layered structure, as evidenced by the dominant cell types on each spatial location, which showed clear segmentation among layers. By contrast, CARD showed a vague boundary between the ONL and GL in the lower left of the tissue section, while RCTD failed to clearly distinguish the ONL from GL and showed a blurry boundary between the GCL and MCL / GL at the top of the tissue section. Meanwhile, iMOSCATO (NB) overestimated the proportions of the cell types GC and OSNs. To further assess performance, we also evaluated the agreement between the manual annotation and dominant cell type. iMOSCATO achieved the highest ARI and NMI values, followed by CARD (see Figure 3C and Table **??**4). Furthermore, each cell type was highly expressed in its corresponding layers, as shown in Figure 3D. iMOSCATO also detected that 62.5% of spots contained all cell types, while 38.5% contained 2, 3, or 4 cell types (see Figures S3A and S3B), supporting our assumption of cell-type sparsity. These results demonstrate that the joint model of iMOSCATO, combined with accounting for cell-type sparsity through the ZIDD model, can enhance cell-type deconvolution performance. We also compared the spatial domain detection results of iMOSCATO with iMOSCATO (NB), BayesSpace, and Louvain, using their default settings, and set the number of domains to four (*D* = 4) for all methods. As shown in Figure 3A, iMOSCATO (ARI = 0.585) achieved the best performance, while iMOSCATO (NB) (ARI = 0.5), BayesSpace (ARI = 0.572), and Louvain (ARI = 0.535) produced inferior results. All methods successfully distinguished the GCL and ONL layers, but compared to iMOSCATO, the other methods produced blurrier boundaries between the MCL and GL layers, leading to compromised performance. In conclusion, the superior performance of iMOSCATO highlights the benefits of integrating both molecular and spatial profiles in the clustering analysis of SRT data, along with the consideration of zero-inflation.

**Figure 3:**
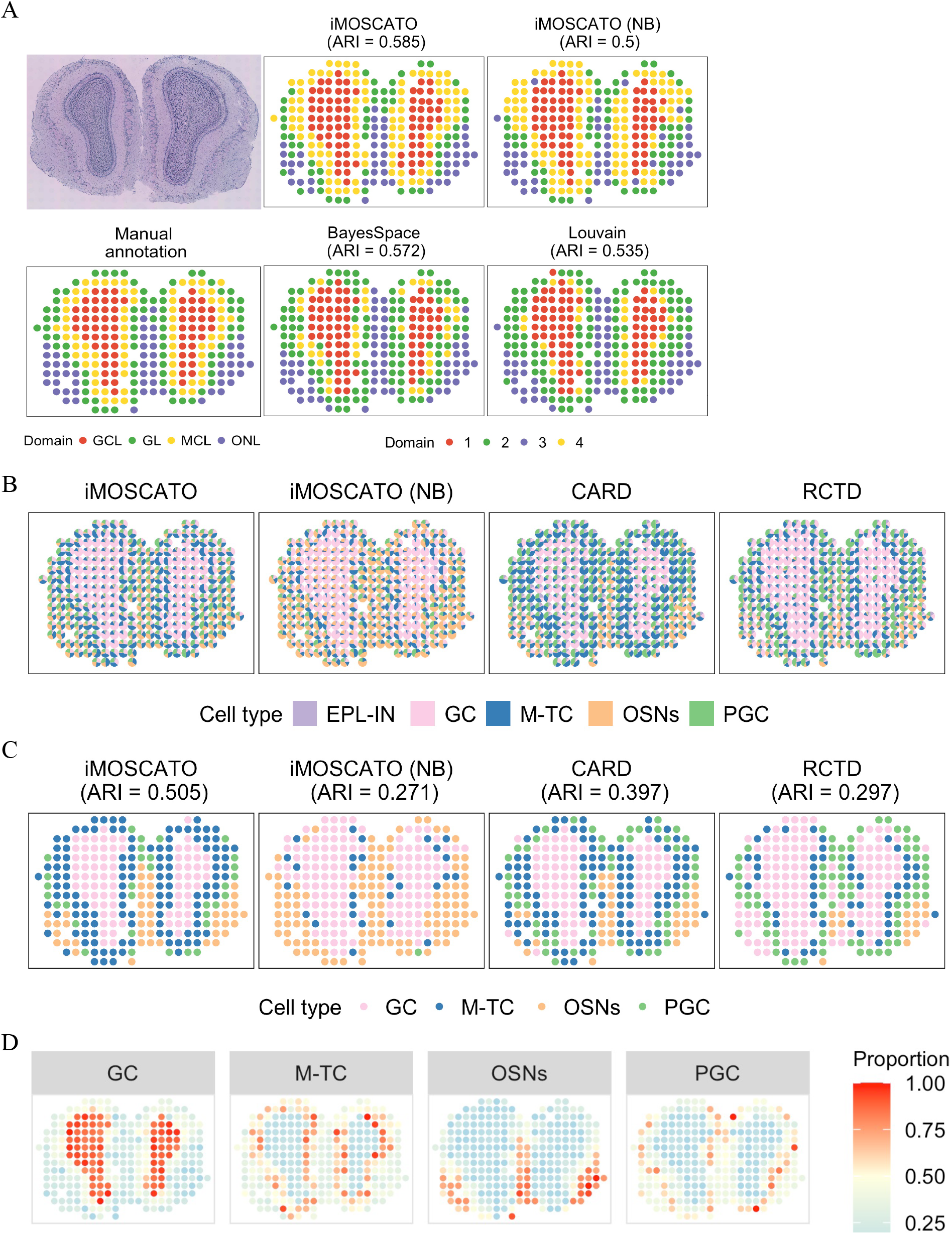
The mouse olfactory bulb ST data analysis. A. The hematoxylin and eosin (H&E)- stained image of the tissue section with manual annotation, and spatial domains detected by different spatial domain detection methods. B. The spatial scatter pie plots of inferred cell-type composition on each spatial location from different cell-type deconvolution methods. C. The dominant cell type on each spatial location from different cell-type deconvolution methods. D. The inferred proportion of each of the four cell types on each spatial location from iMOSCATO.

### 5.2 Application to the Human Pancreatic Ductal Adenocarcinoma ST Data

The second dataset we evaluated was a human pancreatic ductal adenocarcinoma (PDAC) ST data (Moncada et al., 2020). Gene expression was measured from a section of human pancreatic tissue with invasive ductal adenocarcinoma using the ST platform, along with manual annotation for evaluating spatial domain detection performance. This dataset contains three distinct tissue regions (cancer, ductal epithelium, and interstitium) as shown in Figure 4A. For deconvolution, we used a matched scRNA-seq dataset from the same individual obtained from inDrop platform. (Moncada et al., 2020) (denoted as PDAC-B). This scRNA-seq dataset contains 19,736 genes and 1,733 cells, representing 11 cell types. Detailed cell type information is provided in Table S3 in the supplementary materials. We followed the same analysis procedure as described in Section 5.1, but filtered out genes that have zero count on all spots in the SRT data due to its higher zero-inflation. After the quality control procedure, 224 spots with 14,907 genes remained in the SRT data, while the scRNA-seq data retained cells with 15,919 genes. The pairwise PCCs of cell-type proportions ranged from 0.91 to 0.92 across four MCMC chains (Figure S4A), and the trace plot of one cell-type proportion (Figure S4B) further supports good convergence.

**Figure 4:**
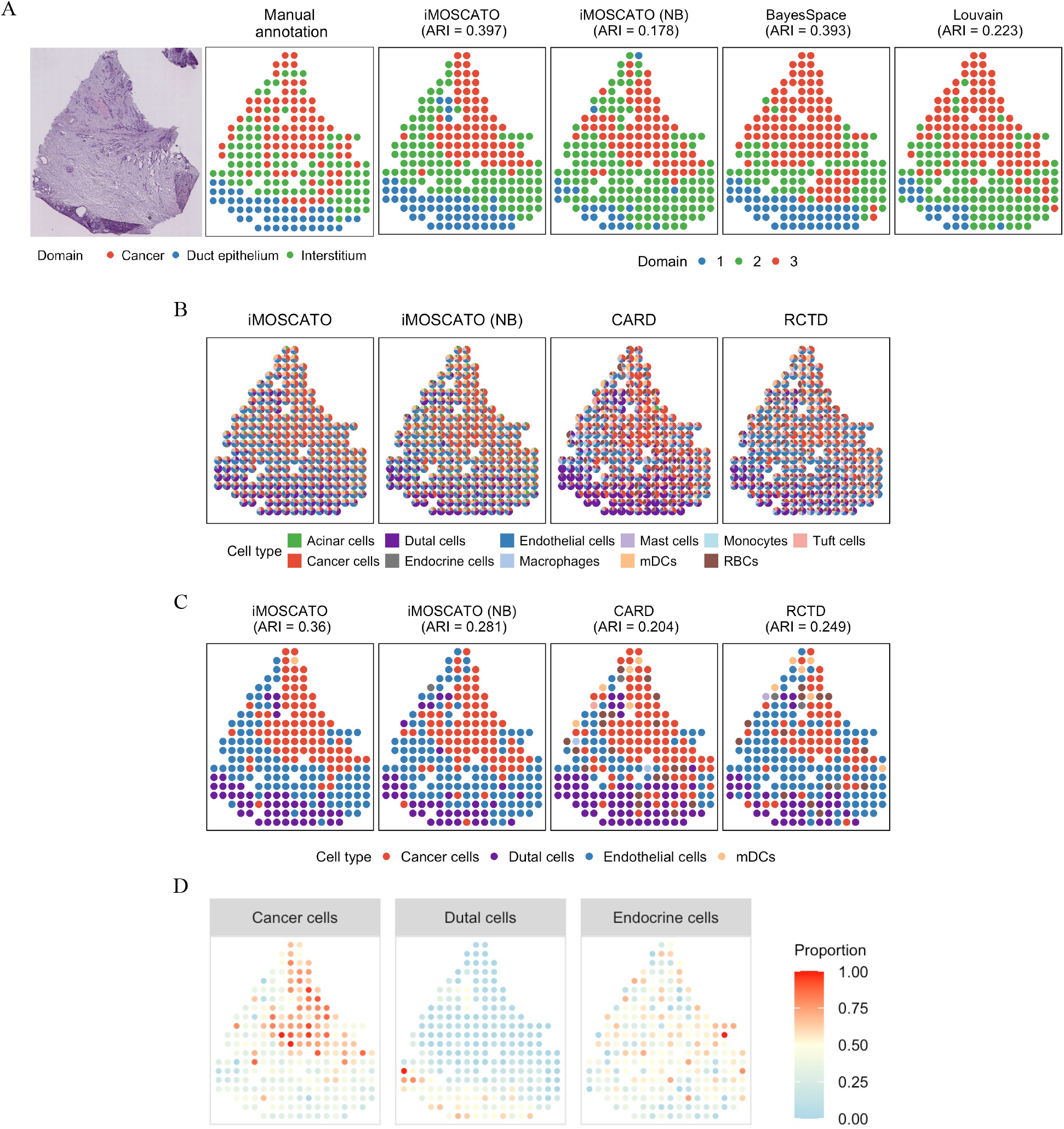
The human pancreatic ductal adenocarcinoma ST data analysis. A. The hematoxylin and eosin (H&E)-stained image of the tissue section with manual annotation, and spatial domains detected by different spatial domain detection methods. B. The spatial scatter pie plots of inferred cell-type composition on each spatial location from different deconvolution methods. C. Dominant cell type on each spatial location from different cell-type deconvolution methods. D. The inferred proportion of each of the three cell types on each spatial location from iMOSCATO.

First, we compared the cell-type compositions detected by iMOSCATO, iMOSCATO (NB), and other competing methods. As shown in Figures 4B and 4C, iMOSCATO captured distinct and well-structured spatial patterns, providing clearer and more biologically consistent distribution of cell types compared to iMOSCATO (NB) and alternative methods. Furthermore, iMOSCATO achieved the highest value of ARI (see Figure 4C), indicating the strongest agreement between the dominant cell types and manual annotations. This highlights its effectiveness in accurately detecting spatial patterns of cell-type composition. In contrast, other methods exhibited more overlapping and ambiguous assignments (see Figures 4C and 4D), demonstrating lower spatial resolution in distinguishing different regions. Among the 224 spots analyzed, 23.6% contained all 11 cell types, while the majority of spots included a number of cell types ranging from 6 to 10 (see Figures S4C and S4D). These findings reinforce the advantage of iMOSCATO in accurately resolving complex cellular heterogeneity and spatial organization in SRT data.

Next, we evaluated the spatial domain detection performance of iMOSCATO. We observed that iMOSCATO achieved the highest consistency with manual annotation, with an ARI of 0.397, followed by BayesSpace (ARI = 0.393). In contrast, iMOSCATO (NB) and Louvain failed to differentiate the duct epithelium and interstitium regions, as shown in Figure 4A. Incorporating spatial information enabled iMOSCATO and BayesSpace to achieve satisfactory clustering performance. Furthermore, the comparison of iMOSCATO (NB) highlights the advantages of modeling zero-inflation in SRT data.

## 6. Discussion

In this paper, we presented iMOSCATO, a fully Bayesian model that integrates scRNA-seq and SRT data for cell-type deconvolution and spatial domain detection. iMOSCATO offers several advantages over existing methods. First, it directly models count data using a NB distribution, which accounts for over-dispersion more effectively than a Poisson distribution. Second, it addresses the challenge of excess zero counts observed in SRT data by incorporating a ZINB model, resulting in more robust performance. Third, it utilizes a ZIDD model to capture the cell-type sparsity existing in cell-type compositions. This approach enables simultaneous deconvolution and spatial domain detection, enhancing overall performance. Moreover, the ZIDD model captures the sparsity inherent in cell-type compositions and provides more interpretable results, particularly in situations where certain cell types are rare or absent in specific regions of interest. Finally, by adopting a Bayesian approach, iMOSCATO improves parameter estimation and quantifies uncertainties, leading to more reliable and precise inferences. In our simulation study, iMOSCATO outperformed all other deconvolution and clustering methods, particularly in datasets with a high proportion of zero counts and proportions. In real data analysis, iMOSCATO demonstrated superior accuracy in deconvolution and clustering by incorporating spatial localization information and utilizing the ZIDD model.

iMOSCATO primarily relies on the SRT molecular profiles, which may limit its ability to distinguish regions that exhibit similar gene expression patterns but differ in their morphological features, as seen in paired histology or pathology images. Studies (Hu et al., 2021; Jiang et al., 2023; Guo et al., 2024) have demonstrated that integrating imaging information significantly enhances the accuracy of spatial domain detection, particularly when manual annotations by pathologists serve as the benchmark for true segmentation. Another limitation is the fixed number of domains *K* in iMOSCATO, although *K* is typically determined by experienced pathologists or through mBIC criterion. To address these limitations, several extensions are worth exploration. Although deconvolution methods infer cell-type proportions in each spot, they cannot estimate the absolute cell-type counts. iMOSCATO could build on approaches from studies (Jiang et al., 2023; Guo et al., 2024) to integrate histology images and predict the total cell count in each spot. Subsequently, a Dirichlet-multinomial distribution can be employed to predict the cell-type counts. Additionally, iMOSCATO could be extended to estimate *K* using a Dirichlet process mixture model (Müller et al., 2015; Li et al., 2017), which would not only estimate *K* but also quantify its uncertainty. As multi-sample analyses become more common, extending the model to handle multiple samples could further enhance the robustness and reliability of clustering results Jiang et al. (2023); Guo et al. (2024). Implementing these extensions, individually or in combination, could yield more precise and reliable spatial deconvolution and clustering outcomes.

## Supporting information

Supplementary Materials for iMOSCATO

## Supplementary Materials

All simulated and real datasets utilized in our analysis, along with the associated source code in R/C++ languages are available at https://github.com/huimin230/iMOSCATO.

## Data Availability

The authors analyzed two publicly available SRT datasets. Mouse olfactory bulb ST data is accessible on the website of the Spatial Research Lab at the KTH Royal Institute of Technology (https://www.spatialresearch.org/). Human pancreatic ductal adenocarcinoma ST data is accessible on the Gene Expression Omnibus (GEO) under accession number GSE111672 (https://www.ncbi.nlm.nih.gov/geo/query/acc.cgi?acc=GSE111672).

