## Supplementary Materials for iMOSCATO for "Robust Bayesian Integrative Modeling of Single Cell and Spatially Resolved Transcriptomics Data"

**by**

**Huimin Li, Bencong Zhu, Xi Jiang, Ying Ma, Lin Xu, and Qiwei Li**

### Web Appendix A. Full Details of the MCMC Algorithm

The model parameter space is $\{\boldsymbol{\Pi}, \boldsymbol{\Omega}, \boldsymbol{\Delta}, \boldsymbol{z}, \boldsymbol{A}, \boldsymbol{H}, \boldsymbol{\phi}\}$, where $\boldsymbol{\Pi} = \{\pi_{ik}, i = 1, \cdots, N, k = 1, \cdots, K\}$ is the cell type proportion matrix, $\boldsymbol{\Omega} = \{\omega_{ik}, i = 1, \cdots, N, k = 1, \cdots, K\}$ is the unnormalized cell type proportion matrix, $\boldsymbol{\Delta} = \{\delta_{ik}, i = 1, \cdots, N, k = 1, \cdots, K\}$ is the extra zero indicator matrix for $\boldsymbol{\Omega}$, $\boldsymbol{z} = \{z_i, i = 1, \cdots, N\}$ is the spatial domain allocation vector, $\boldsymbol{A} = \{\alpha_{dk}, d = 1, \cdots, D, k = 1, \cdots, K\}$ is the concentration parameter matrix, $\boldsymbol{H} = \{\eta_{ij}, i = 1, \cdots, N, j = 1, \cdots, P\}$ is the extra zero indicator matrix for $\boldsymbol{Y}$, and $\boldsymbol{\phi} = \{\phi_j, j = 1, \cdots, P\}$ is the collection of dispersion parameters of spatial transcriptomics (ST) data.

We start by writing the full posterior,

$$p(\boldsymbol{\Pi}, \boldsymbol{\Omega}, \boldsymbol{\Delta}, \boldsymbol{z}, \boldsymbol{A}, \boldsymbol{H}, \boldsymbol{\phi}|\boldsymbol{Y}) \quad \propto \quad L(\boldsymbol{Y}|\boldsymbol{\Pi}, \boldsymbol{H}, \boldsymbol{\phi})p(\boldsymbol{\Omega}|\boldsymbol{z}, \boldsymbol{A}, \boldsymbol{\Delta})p(\boldsymbol{\Delta})p(\boldsymbol{A})p(\boldsymbol{z})p(\boldsymbol{H})p(\boldsymbol{\phi}).$$

According to Section 2.2, we can compute the data likelihood and priors as

$$\begin{aligned}
L(\boldsymbol{Y}|\boldsymbol{\Pi}, \boldsymbol{H}, \boldsymbol{\phi}) &= \prod_{i=1}^{N} \prod_{\{j:\eta_{ij}=0\}} \mathrm{NB}\left(y_{ij}; s_i \sum_{k=1}^{K} \pi_{ik} b_{kj}, \phi_j\right). \\
\pi(\boldsymbol{H}) &= \prod_{i=1}^{N} \prod_{j=1}^{P} \mathrm{Bern}(\eta_{ij}; \theta). \\
\pi(\boldsymbol{\phi}) &= \prod_{j=1}^{P} \mathrm{Ga}(\phi_j; a_\phi, b_\phi).
\end{aligned}$$

We can also obtain the likelihood of spot $i$ and gene $j$ by

$$\begin{aligned}
L(\boldsymbol{y}_{i\cdot}|\boldsymbol{\pi}_{i\cdot}, \boldsymbol{\eta}_{i\cdot}, \boldsymbol{\phi}) &= \prod_{\{j:\eta_{ij}=0\}} \mathrm{NB}\left(y_{ij}; s_i \sum_{k=1}^{K} \pi_{ik} b_{kj}, \phi_j\right). \\
L(\boldsymbol{y}_{\cdot j}|\boldsymbol{\Pi}, \boldsymbol{\eta}_{\cdot j}, \phi_j) &= \prod_{\{i:\eta_{ij}=0\}} \mathrm{NB}\left(y_{ij}; s_i \sum_{k=1}^{K} \pi_{ik} b_{kj}, \phi_j\right).
\end{aligned}$$

According to Sections 2.3 and 2.4, we can calculate the rest of priors as

$$\begin{aligned}
p(\boldsymbol{\Omega}|\boldsymbol{z}, \boldsymbol{A}, \boldsymbol{\Delta}) &= \prod_{i=1}^{N} \prod_{\{k:\delta_{ik}=0\}} \mathrm{Ga}(\omega_{ik}; \alpha_{dk}, 1). \\
p(\boldsymbol{\Delta}) &= \prod_{i=1}^{N} \prod_{k=1}^{K} \mathrm{Bern}(\delta_{ik}; q).
\end{aligned}$$

$$p(\boldsymbol{A}) = \prod_{d=1}^{D} \prod_{\{k:\delta_{ik}=0\}} \text{Ga}(\alpha_{dk}; a_\alpha, b_\alpha).$$

$$\pi(\boldsymbol{z}) \propto \exp\left\{\sum_{d=1}^{D} e_d \sum_{i=1}^{n} \text{I}(z_i = d) + h \sum_{i<i'} g_{ii'} \text{I}(z_{i'} = z_i)\right\}.$$

The pmf's or pdf's of the involved common distributions are given below:

$$\text{If} \quad x \sim \text{NB}(\mu, \phi), \quad \text{then} \quad p(x) = \frac{\Gamma(x+\phi)}{x!\Gamma(\phi)} \left(\frac{\phi}{\mu+\phi}\right)^{\phi} \left(\frac{\mu}{\mu+\phi}\right)^{x}.$$

$$\text{If} \quad \boldsymbol{x} \sim \text{Dir}(\boldsymbol{\alpha}), \quad \text{then} \quad p(x_1, \cdots, x_K | \alpha_1, \cdots, \alpha_K) = \frac{\Gamma(\sum_{k=1}^{K} \alpha_k)}{\prod_{k=1}^{K} \Gamma(\alpha_k)} \prod_{k=1}^{K} x_k^{\alpha_k - 1}.$$

$$\text{If} \quad x \sim \text{Bern}(\theta), \quad \text{then} \quad p(x) = \theta^x (1-\theta)^{1-x}.$$

$$\text{If} \quad x \sim \text{Ga}(a, b), \quad \text{then} \quad p(x) = \frac{b^a}{\Gamma(a)} x^{a-1} \exp(-bx).$$

We detail the MCMC algorithm below. At each MCMC iteration, We perform the following steps:

**Joint update of zero-inflation parameter $\boldsymbol{\Delta}$, cell type proportion $\boldsymbol{\Pi}$, and unnormalized cell type proportion $\boldsymbol{\Omega}$:** Since $\pi_{ik} = \omega_{ik} / \sum_{k'=1}^{K} \omega_{ik'}$, updating of $\boldsymbol{\Omega}$ is equivalent to update $\boldsymbol{\Pi}$. We perform a between-model step to update these parameters jointly since $\boldsymbol{\Pi}$ and $\boldsymbol{\Omega}$ depends on $\boldsymbol{\Delta}$. This is done via an *add-delete* algorithm. In this approach, a new candidate vector $\boldsymbol{\delta}_{i\cdot}^*$, $i = 1, \cdots, N$, is generated by randomly choosing an entry of $\boldsymbol{\delta}_{i\cdot}$, say $k$, $k = 1, \cdots, K$, and change its value to $\delta_{ik}^* = 1 - \delta_{ik}$. Then, this proposed move is accepted with probability min(1, $\text{m}_{\text{MH}}$), where the Hastings ratio is

$$\begin{aligned}
\text{m}_{\text{MH}} &= \frac{f(\boldsymbol{y}_{i\cdot}|\boldsymbol{\pi}_{i\cdot}^*, \boldsymbol{\eta}_{i\cdot}, \boldsymbol{\phi})}{f(\boldsymbol{y}_{i\cdot}|\boldsymbol{\pi}_{i\cdot}, \boldsymbol{\eta}_{i\cdot}, \boldsymbol{\phi})} \frac{p(\omega_{ik}^*|z_i, \alpha_{dk}, \delta_{ik}^*)}{p(\omega_{ik}|z_i, \alpha_{dk}, \delta_{ik})} \frac{p(\delta_{ik}^*)}{p(\delta_{ik})} \frac{J(\omega_{ik} \leftarrow \omega_{ik}^*|\delta_{ik} \leftarrow \delta_{ik}^*)}{J(\omega_{ik}^* \leftarrow \omega_{ik}|\delta_{ik}^* \leftarrow \delta_{ik})} \frac{J(\delta_{ik} \leftarrow \delta_{ik}^*)}{J(\delta_{ik}^* \leftarrow \delta_{ik})} \\
&= \frac{\prod_{\{j:\eta_{ij}=0\}} \text{NB}\left(y_{ij}; s_i \sum_{k=1}^{K} \pi_{ik}^* b_{kj}, \phi_j\right)}{\prod_{\{j:\eta_{ij}=0\}} \text{NB}\left(y_{ij}; s_i \sum_{k=1}^{K} \pi_{ik} b_{kj}, \phi_j\right)} \frac{(\text{Ga}(\omega_{ik}^*; \alpha_{dk}, 1))^{1-\delta_{ik}^*}}{(\text{Ga}(\omega_{ik}; \alpha_{dk}, 1))^{1-\delta_{ik}}} \frac{\text{Bern}(\delta_{ik}^*; q)}{\text{Bern}(\delta_{ik}; q)} \\
&\quad \frac{J(\omega_{ik} \leftarrow \omega_{ik}^*|\delta_{ik} \leftarrow \delta_{ik}^*)}{J(\omega_{ik}^* \leftarrow \omega_{ik}|\delta_{ik}^* \leftarrow \delta_{ik})}.
\end{aligned}$$

while the last proposal density ratio $J(\delta_{ik} \leftarrow \delta_{ik}^*)/J(\delta_{ik}^* \leftarrow \delta_{ik})$ equals one. We use $J(\cdot \leftarrow \cdot)$ to denote the proposal probability function for selected move. For the *add* step (i.e., $\delta_{ik}^* = 1$),

$\omega^*_{ik} = 0$. For the *delete* step (i.e., $\delta^*_{ik} = 0$), we propose a new $\omega^*_{ik}$ from $\mathrm{Ga}(\alpha_{dk}, 1)$. Then,

$$\frac{J(\omega_{ik} \leftarrow \omega^*_{ik} | \delta_{ik} \leftarrow \delta^*_{ik})}{J(\omega^*_{ik} \leftarrow \omega_{ik} | \delta^*_{ik} \leftarrow \delta_{ik})} = \begin{cases} \mathrm{Ga}(\omega_{ik}; \alpha_{dk}, 1) & \text{for } \textit{add step} \\ 1/\mathrm{Ga}(\omega^*_{ik}; \alpha_{dk}, 1) & \text{for } \textit{delete} \text{ step} \end{cases}.$$

**Update of the cell type proportion $\mathbf{\Pi}$ and unnormalized cell type proportion $\mathbf{\Omega}$:** Since $\pi_{ik} = \omega_{ik} / \sum_{k'=1}^{K} \omega_{ik'}$, updating of $\omega_{ik}$ is equivalent to update $\pi_{ik}$. We perform a random walk Metropolis-Hastings (RWMH) algorithm to update each $\omega_{ik}$, $i = 1, \cdots, N$, $k = 1, \cdots, K$ sequentially only for those $\delta_{ik} = 0$. Particularly, we propose a new $\log \omega^*_{ik}$ from $\mathrm{N}(\log \omega_{ik}, \tau^2_\omega)$ and then accept the proposed value $\omega^*_{ik}$ with probability $\min(1, \mathrm{m}_{\mathrm{MH}})$, where

$$\mathrm{m}_{\mathrm{MH}} = \frac{\prod_{\{j:\eta_{ij}=0\}} \mathrm{NB}\left(y_{ij}; s_i \sum_{k=1}^{K} \pi^*_{ik} b_{kj}, \phi_j\right)}{\prod_{\{j:\eta_{ij}=0\}} \mathrm{NB}\left(y_{ij}; s_i \sum_{k=1}^{K} \pi_{ik} b_{kj}, \phi_j\right)} \frac{\mathrm{Ga}(\omega^*_{ik}; \alpha_{dk}, 1)}{\mathrm{Ga}(\omega_{ik}; \alpha_{dk}, 1)} \frac{J(\omega_{ik} \leftarrow \omega^*_{ik})}{J(\omega^*_{ik} \leftarrow \omega_{ik})}$$

Note that the proposal density ratio equals one for this RWMH update.

**Update of the spatial domain allocation $\boldsymbol{z}$:** We update each $z_i$, $i = 1, \cdots, N$ separately using a Gibbs Sampler. At iteration, we draw a new value $z_i = d$ with the probability $p(z_i = d|\cdot) / \sum_{d'=1}^{D} p(z_i = d'|\cdot)$, where

$$p(z_i = d|\cdot) \propto \left\{ \prod_{\{k:\delta_{ik}=0\}} \mathrm{Ga}(\omega_{ik}; \alpha_{dk}, 1) \right\} \exp\left\{ e_d + h \sum_{i'=1}^{N} g_{ii'} \mathrm{I}(z_{i'} = z_i) \right\}$$

**Update of the concentration parameter $\boldsymbol{A}$:** We update each $\alpha_{dk}$, $i = 1, \cdots, N, k = 1, \cdots, K$ separately via the RWMH algorithm. We first propose a new $\log \alpha^*_{dk}$ from $\mathrm{N}(\log \alpha_{dk}, \tau^2_\alpha)$, and then accept the proposed value with probability $\min(1,\ \mathrm{m}_{\mathrm{MH}})$, where

$$\mathrm{m}_{\mathrm{MH}} = \frac{\prod_{\{i:z_i=d,\delta_{ik}=0\}} \mathrm{Ga}(\omega_{ik}; \alpha^*_{dk}, 1)}{\prod_{\{i:z_i=d,\delta_{ik}=0\}} \mathrm{Ga}(\omega_{ik}; \alpha_{dk}, 1)} \frac{\mathrm{Ga}(\alpha^*_{dk}; a_\alpha, b_\alpha)}{\mathrm{Ga}(\alpha_{dk}; a_\alpha, b_\alpha)} \frac{J(\alpha_{ik} \leftarrow \alpha^*_{ik})}{J(\alpha^*_{ik} \leftarrow \alpha_{ik})}$$

Note that the proposal density ratio equals one for this RWMH update.

**Update of the zero-inflation parameter $\boldsymbol{H}$:** We update each false zero indicator $\eta_{ij}$, $i = 1, \cdots, N$, $j = 1, \cdots, P$ that corresponds to $y_{ij} = 0$ using the Gibbs sampler,

$$p(\eta_{ij}|\cdot) \propto \mathrm{NB}\left(y_{ij}; s_i \sum_{k=1}^{K} \pi_{ik} b_{kj}, \phi_j\right) \mathrm{Bern}(\eta_{ij}; \theta)$$

$$\eta_{ij}|\cdot \sim \mathrm{Bern}\left(\frac{p(\eta_{ij} = 1|\cdot)}{p(\eta_{ij} = 1|\cdot) + p(\eta_{ij} = 0|\cdot)}\right)$$

**Update of the dispersion parameter $\phi$:** We perform a RWMH algorithm to update each $\phi_j, j = 1, \cdots, P$ sequentially. Particularly, we propose a new we propose a new $\log \phi_j^*$ from $\mathrm{N}\left(\log \phi_j, \tau_\phi^2\right)$ and then accept the proposed value $\phi_j^*$ with probability $\min(1, \mathrm{m}_{\mathrm{MH}})$, where

$$\mathrm{m}_{\mathrm{MH}} = \frac{\prod_{\{i:\eta_{ij}=0\}} \mathrm{NB}\left(y_{ij}; s_i \sum_{k=1}^K \pi_{ik} b_{kj}, \phi_j^*\right)}{\prod_{\{i:\eta_{ij}=0\}} \mathrm{NB}\left(y_{ij}; s_i \sum_{k=1}^K \pi_{ik} b_{kj}, \phi_j\right)} \frac{\mathrm{Ga}(\phi_j^*; a_\phi, b_\phi)}{\mathrm{Ga}(\phi_j; a_\phi, b_\phi)} \frac{J(\phi_j \leftarrow \phi_j^*)}{J(\phi_j^* \leftarrow \phi_j)}$$

Note that the proposal density ratio equals one for this RWMH update.

### Web Appendix B. Supplement to Section 4: Simulation Study

Web Appendix B.1 *Data Generative Model*

For the cell-level count data, we created $M = 500$ cells with $K = 4$ cell types, where cell types 1, 2, 3, and 4 have 200, 50, 100, and 150 cells, respectively. To characterize the excess zeros and over-dispersion in the count data, we simulated the observed gene expression counts $x_{mj}$ from a zero-inflated Negative binomial (ZINB) mixture model,

$$x_{mj} \sim \vartheta_m \mathrm{I}(x_{mj} = 0) + (1 - \vartheta_m)\mathrm{NB}(v_m b_{kj}, \psi_j)$$

The size factor $v_m$ were independent and identically distributed (i.i.d.) from LN(0, $0.2^2$) and the gene-specific dispersion parameter $\psi_j$ were i.i.d from Exp(0.1) with mean 10. For the choice of extra zero proportion $\vartheta_m$, we randomly selected 5%, 10%, and 30% counts and manually set their values to zero.

For each gene $j$ at cell type $k$, its latent normalized expression level was generated as:

$$\log(b_{kj}) = \begin{cases} \beta_0 + d_k + \varepsilon_j & \text{if gene } j \text{ is a cell-type marker gene} \\ \beta_0 + \varepsilon_j & \text{if gene } j \text{ is a non-cell-type marker gene} \end{cases}$$

where $\beta_0$ is the baseline of log-normalized expression level and $\varepsilon_j$ is the gene-specific non-spatial random error following $\varepsilon_j \sim \mathrm{N}(0, \sigma_j^2)$, and $d_k$ is cell type-specific fold change between informative and non-cell-type marker genes. We set $\beta_0 = 2$, $\sigma_j \sim \mathrm{Exp}(0.3)$, $d_1, d_2, d_3$, and $d_4$ are randomly chosen from $0, \log 3, \log 6$, and $\log 18$ without replacement for each cell-type marker gene $j$. For a non-cell-type marker gene, the normalized expression levels are i.i.d. from a log-normal (LN) distribution with mean 2 and variance $\sigma_j^2$.

In the simulation of spot-level data, 260 spots were assigned to two spatial domains. We assumed that each domain is characterized by a unique dominant cell type, with cell types 1 and 4 serving as the dominant cell types for spatial domains 1 and 2, respectively. The unnormalized cell type proportion $\omega_{ik}$ were i.i.d from a zero-inflated Gamma distribution as follows:

$$\omega_{ik} \quad \sim \quad q_i \mathrm{I}(\omega_{ik} = 0) + (1 - q_i)\mathrm{Ga}(\alpha_{dk}, 1)$$

where the concentration parameters $\boldsymbol{\alpha}_{d\cdot} = (3, 1, 1, 1)$ and $(1, 1, 1, 3)$ for spatial domains 1 and 2 respectively. For the choice of an extra zero proportion $q_i$, we randomly selected 10%, 30%, and 50% non-dominant cell type proportions and manually forced their values to zero. Then the proportion of cell type $\pi_{ik}$ is $\omega_{ik} / \sum_{k'=1}^{K} \omega_{i'k}$. To characterize the excess zeros and over-dispersion in the count data, we simulated the expression count $y_{ij}$ from a ZINB distribution as follows:

$$y_{ij} \quad \sim \quad \theta_i \mathrm{I}(y_{ij} = 0) + (1 - \theta_i)\mathrm{NB}\left(s_i \sum_{k=1}^{K} \pi_{ik} b_{kj}, \phi_j\right)$$

The size factor $s_i$ were i.i.d. from LN(0, $0.2^2$) and the gene-specific dispersion parameter $\phi_j$ were i.i.d from Exp(0.1). For the choice of extra zero proportion $\theta_i$, we randomly selected 5%, 10%, and 30% counts and manually set their values to zero. Combined with three zero-inflation and three cell type sparsity settings, there were nine different scenarios in total. For each scenario, we independently repeated the above steps to generate 30 replicates.

#### Web Appendix B.2 *Prior and Algorithm Settings*

For the hyperparameter settings, we used the following defaults. We set the hyperparameters $\theta$ and $q$ that control the proportion of extra zeros as 0.5. For the gamma priors on the negative binomial (NB) dispersion parameters and concentration parameters, i.e., $\phi_j \sim \mathrm{Ga}(a_\phi, b_\phi)$ and

$\alpha_{dk} \sim \text{Ga}(a_\alpha, b_\alpha)$, we set $a_\phi, b_\phi, a_\alpha$ and $b_\alpha$ to 0.001, resulting in a vague distribution with mean of 1 and variance of 1,000. For the MRF prior, we set $e_1 = e_2 = 1$ and $h = 1$. The results we report in simulation study were obtained by running one MCMC chain with $5,000$ iterations, discarding the first 50% of sweeps as burn-in. All experiments were implemented in `R` with the `Rcpp` package to accelerate computations.

[Figure 1 about here.]

[Figure 2 about here.]

[Figure 3 about here.]

[Figure 4 about here.]

[Table 1 about here.]

[Table 2 about here.]

[Table 3 about here.]

[Table 4 about here.]

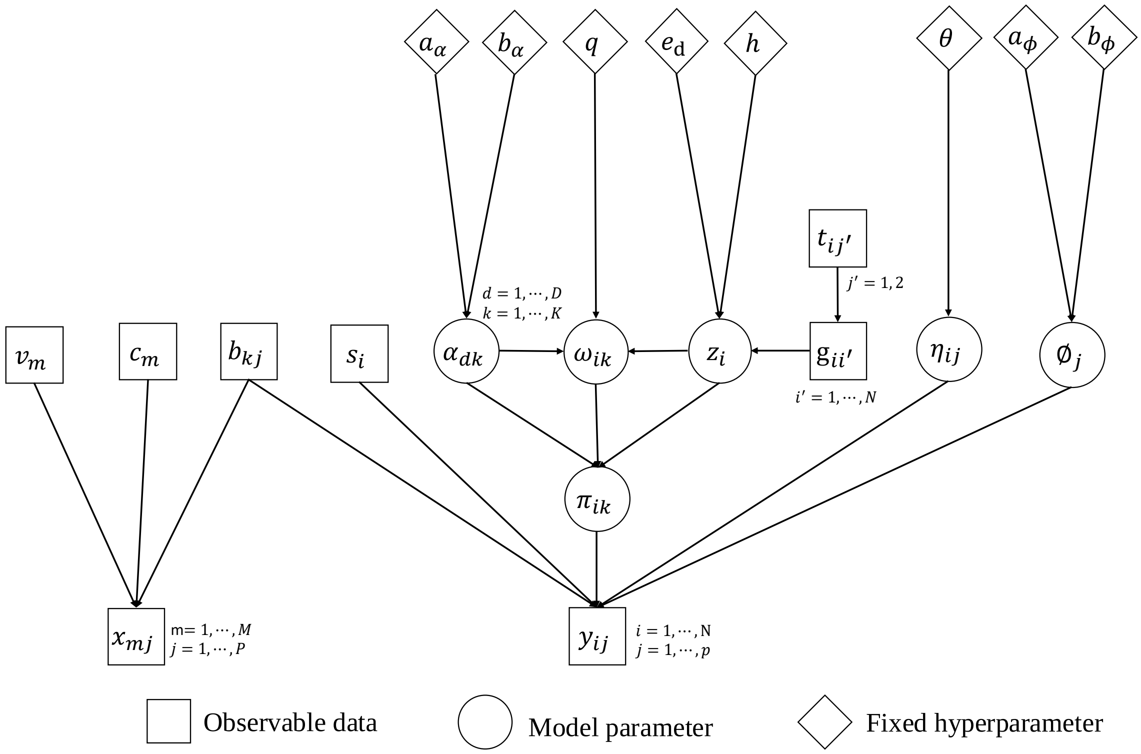


**Figure S1**: Graphical representation of the proposed iMOSCATO model. Square, circle, and diamond-shaped nodes refer to observable data, model parameter, and fixed hyperparameter, respectively. Circles with a dashed outline indicate nuisance parameters that are integrated out in iMOSCATO. A link between two nodes represents a direct probabilistic dependence.

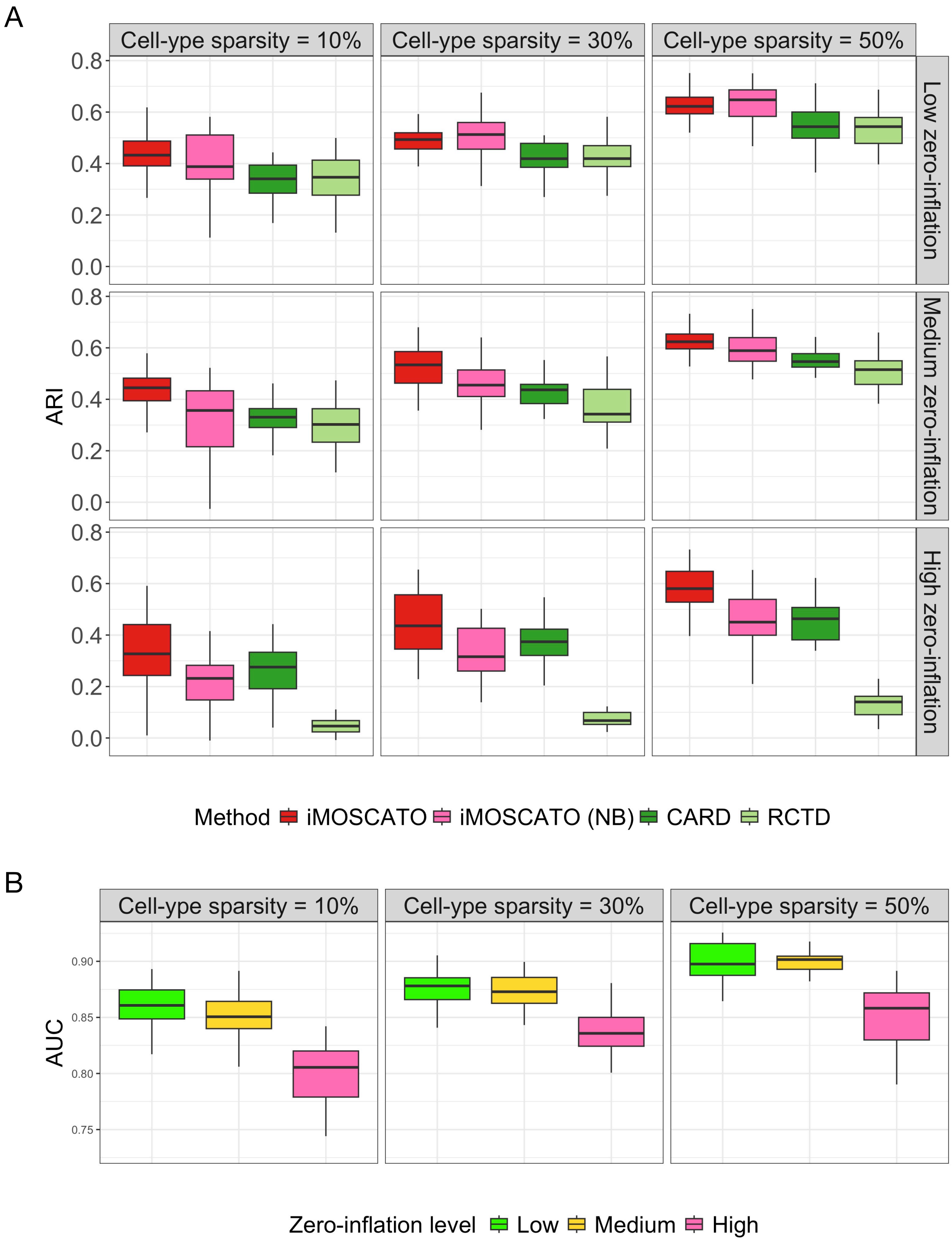


**Figure S2**: The simulation study. A. The boxplots of adjusted Rand indices (ARIs) achieved by different cell-type deconvolution methods under different scenarios in terms of count zero-inflation and cell-type sparsity settings. B. The boxplots of area under curves (AUCs) achieved by iMOSCATO under different scenarios in terms of count zero-inflation and cell-type sparsity settings.

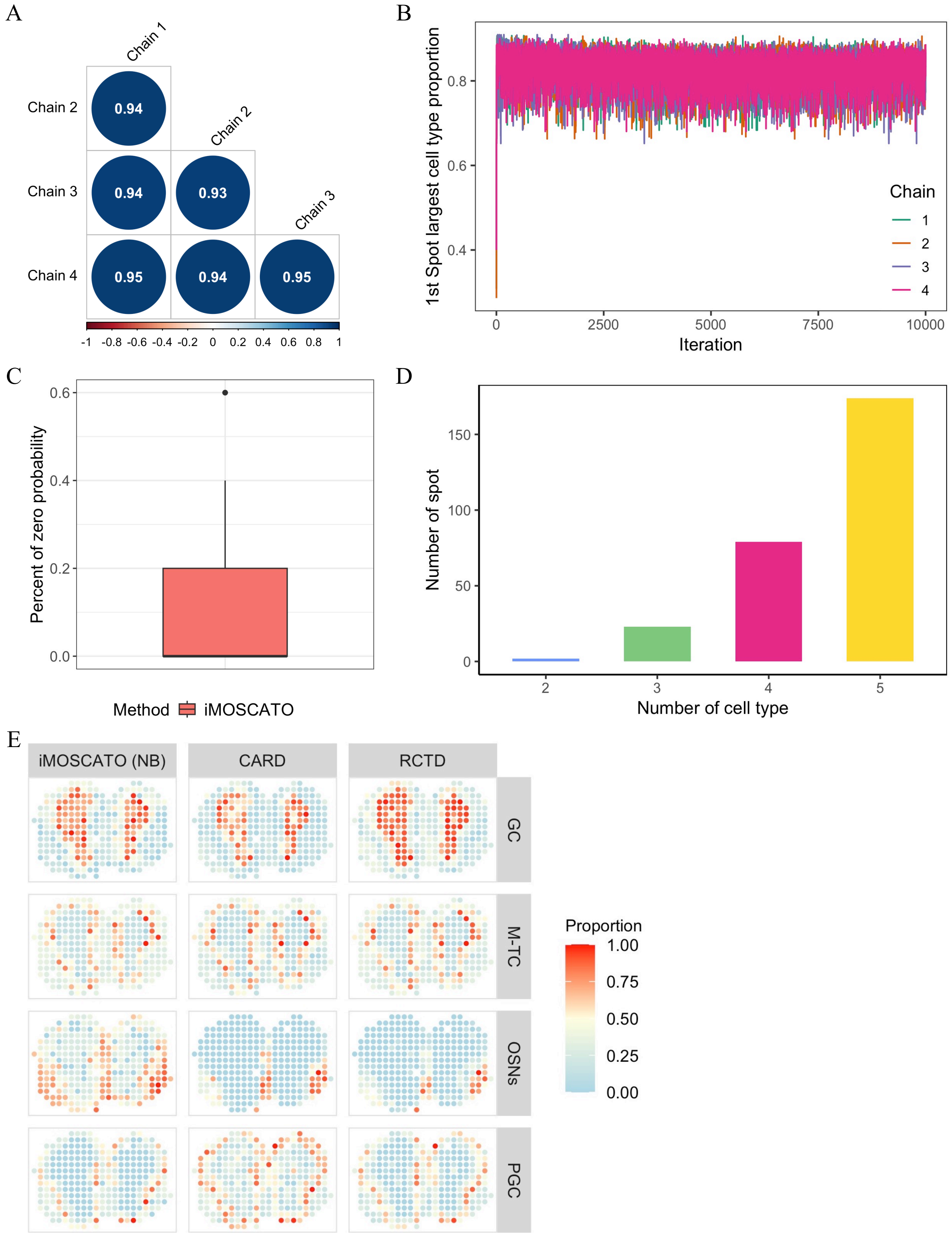


**Figure S3**: The mouse olfactory bulb ST data analysis. A. The plot of pairwise Pearson correlation coefficients of the first spot largest cell type proportion between four independent Markov chains. B. The trace plot of the first spot largest cell type proportion of four independent Markov chains. C. The boxplot of percentage of zero probability on each spatial location achieved by iMOSCATO. D. The bar plot of number of spots with different number of cell types achieved by iMOSCATO. E. The inferred proportion of each of the four cell types on each spatial location from different cell-type deconvolution methods.

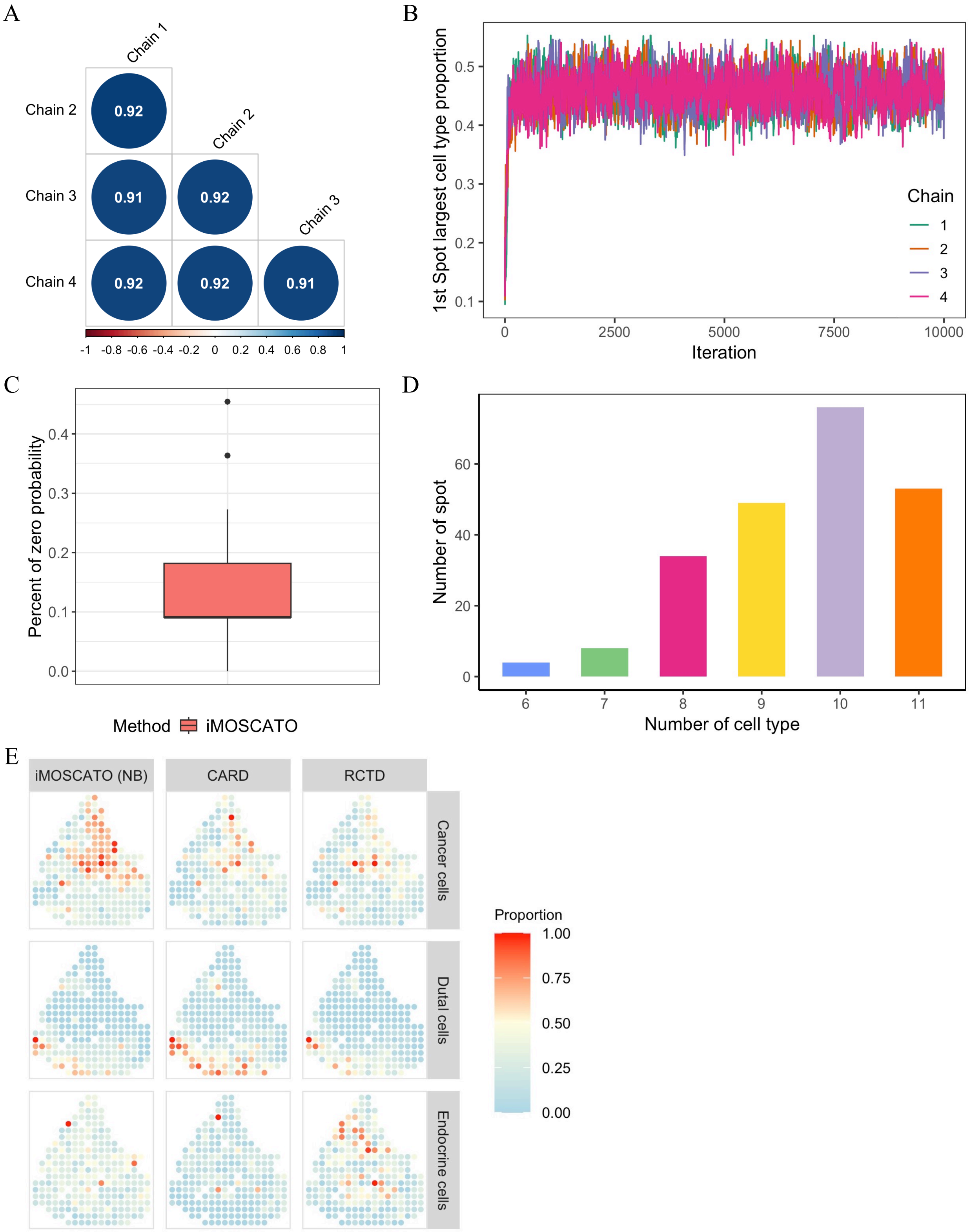


**Figure S4**: The human pancreatic ductal adenocarcinoma ST data analysis. A. The plot of pairwise Pearson correlation coefficients of the first spot largest cell type proportion between four independent Markov chains. B. The trace plot of the first spot largest cell type proportion of four independent Markov chains. C. The boxplot of percentage of zero probability on each spatial location achieved by iMOSCATO. D. The bar plot of number of spots with different number of cell types achieved by iMOSCATO. E. The inferred proportion of each of the three cell types on each spatial location from different cell-type deconvolution methods.

Table S1: Hierarchical formulation of the proposed iMOSCATO model.

**Hierarchical model**:

$$y_{ij}|\pi_{ik}, \eta_{ij}, \phi_j \sim \begin{cases} 0 & \text{if } \eta_{ij} = 1 \\ \text{NB}(y_{ij}; s_i \sum_{k=1}^{K} \pi_{ik} b_{kj}, \phi_j) & \text{if } \eta_{ij} = 0 \end{cases}$$

**Cell type proportion prior**:

$$\boldsymbol{\pi}_{i\cdot}|z_i = d, \boldsymbol{\alpha}_{d\cdot} \sim \text{Dir}(\boldsymbol{\pi}_{i\cdot}; \boldsymbol{\alpha}_{d\cdot})$$

**Unnormalized cell type proportion prior**:

$$\omega_{ik}|z_i = d, \delta_{ik}, \alpha_{dk} \sim \delta_{ik}\text{I}(\omega_{ik} = 0) + (1 - \delta_{ik})\text{Ga}(\alpha_{dk}, 1), \qquad \pi_{ik} = \frac{\omega_{ik}}{\sum_{k'=1}^{K} \omega_{ik'}}$$

**Zero proportion indicator prior**:

$$\delta_{ik}|q \sim \text{Bern}(\delta_{ik}; q)$$

**Concentration parameter prior**:

$$\alpha_{dk} \propto \text{Ga}(\alpha_{dk}; a_\alpha, b_\alpha)$$

**False zero indicator prior**:

$$\eta_{ij}|\theta \sim \text{Bern}(\delta_{ik}; \theta)$$

**Spatial domain Markov random field prior**:

$$p(z_i = d|\boldsymbol{z}_{-i}) \propto \exp\left(e_d + h \sum_{i'=1}^{N} g_{ii'}\text{I}(z_{i'} = d)\right)$$

**Negative binomial dispersion prior**:

$$\phi_j \sim \text{Ga}(\phi_j; a_\phi, b_\phi)$$

**Fixed hyperparameters**:

$$q, a_\alpha, b_\alpha, \theta, e_1, \cdots, e_D, h, a_\phi, b_\phi$$

Table S2: A summary of the two real datasets analyzed in the paper.

| Dataset | Data | No. of samples | No. of genes | No. of clusters | Zero proportion | Reference |
|---|---|---|---|---|---|---|
| Mouse olfactory bulb | scRNA-seq data | 12,801 | 18,506 | 5 | 62% | (Tepe et al., 2018) |
| ST data | ST data | 282 | 16,034 | 4 | 72% | (Ståhl et al., 2016) |
| Human pancreatic ductal | scRNA-seq data | 1,733 | 19,736 | 11 | 93% | (Moncada et al., 2020) |
| adenocarcinoma ST data | ST data | 224 | 19,736 | 3 | 94% | (Moncada et al., 2020) |

Table S3: A Summary of cell type information in scRNA-seq data and spatial domain information in ST data of the two real datasets analyzed in the paper.

| Dataset | Data | Cluster | No. of cells (%) | Cluster | No. of cells (%) |
|---|---|---|---|---|---|
| Mouse olfactory bulb ST data | scRNA-seq data | EPL-IN | 161 (1.3%) | GC | 8,614 (67.3%) |
| | | M-TC | 1,133 (8.9%) | OSNs | 1,200 (9.4%) |
| | | PGC | 1,693 (13.2%) | Total | 1,2801 (100%) |
| | ST data | GCL | 67 (24.2%) | GL | 80 (28.9%) |
| | | MCL | 75 (27.1%) | ONL | 55 (19.9%) |
| | | Total | 277 (100%) | | |
| Human pancreatic ductal adenocarcinoma ST data | scRNA-seq data | Acinar cells | 6.0 (0.3%) | Cancer cells | 339 (19.6%) |
| | | Ductal cells | 1,099 (63.5%) | Endocrine cells | 13 (0.8%) |
| | | Endothelial cells | 159 (9.2%) | Macrophages | 9 (0.5%) |
| | | Mast cells | 13 (0.8%) | mDCs | 35 (2.0%) |
| | | Monocytes | 20 (1.2%) | RBCs | 3 (0.2%) |
| | | Tuft cells | 37 (2.1%) | Total | 1,733 (100.0%) |
| | ST data | Cancer | 88 (39.3%) | Duct Epithelium | 47 (21.0%) |
| | | Interstitium | 89 (39.7%) | Total | 224 (100.0%) |

Table S4: A summary of model performance in terms of ARI and NMI, and computational efficiency in terms of running time (in seconds) of iMOSCATO (per $1,000$ MCMC iterations), CARD, RCTD, BayesSpace, and Louvain on the two real datasets analyzed in the paper. All experiments were conducted on a laptop equipped with an Apple M1 Pro chip and 16GB of memory.

| Dataset | Analysis | Method | ARI | NMI | $P$ | $N$ | Running time |
|---|---|---|---|---|---|---|---|
| Mouse olfactory bulb ST data | Cell-type Deconvolution | iMOSCATO | **0.505** | **0.593** | 412 | 282 | 127.96s |
| | | CARD | 0.397 | 0.470 | 1,991 | | 53.60s |
| | | RCTD | 0.297 | 0.371 | 2,740 | | 180.79s |
| | Spatial domain detection | iMOSCATO | **0.585** | **0.668** | 412 | 282 | 127.96s |
| | | BayesSpace | 0.572 | 0.633 | 15 | | 99.03s |
| | | Louvain | 0.535 | 0.603 | 10 | | 6.92s |
| Human pancreatic ductal adenocarcinoma ST data | Cell-type Deconvolution | iMOSCATO | **0.360** | **0.365** | 1,853 | 224 | 608.82s |
| | | CARD | 0.204 | 0.215 | 6,197 | | 3.93s |
| | | RCTD | 0.249 | 0.230 | 4,955 | | 58.84s |
| | Spatial domain detection | iMOSCATO | **0.397** | 0.423 | 1,853 | 224 | 608.82s |
| | | BayesSpace | 0.393 | **0.458** | 15 | | 56.21s |
| | | Louvain | 0.223 | 0.294 | 50 | | 2.94s |
